# Encoding opposing valences through frequency-dependent transmitter switching in single peptidergic neurons

**DOI:** 10.1101/2024.11.09.622790

**Authors:** Dong-Il Kim, Sukjae J. Kang, Jinho Jhang, Yong S. Jo, Seahyung Park, Mao Ye, Gyeong Hee Pyeon, Geun-Ho Im, Seong-Gi Kim, Sung Han

## Abstract

Peptidergic neurons often co-express fast transmitters and neuropeptides in separate vesicles with distinct release properties. However, the release dynamics of each transmitter in various contexts have not been fully understood in behaving animals. Here, we demonstrate that calcitonin gene-related peptide (CGRP) neurons in the external lateral subdivision of the parabrachial nucleus (CGRP^PBel^) encode opposing valence via differential release, rather than corelease, of glutamate and neuropeptides, according to firing rate. Glutamate is released preferentially at lower firing rates with minimal release at higher firing rates, whereas neuropeptides are released at higher firing rates, resulting in frequency-dependent switching of transmitters. Aversive stimuli evoke high frequency responses with accompanying neuropeptide release to encode negative valence, whereas appetitive stimuli evoke low frequency responses with glutamate release to encode positive valence. Our study reveals a previously unknown capability of single CGRP^PBel^ neurons to bidirectionally encode valence via frequency-dependent differential release of transmitters *in vivo*.

## Introduction

Neuromodulators, such as neuropeptides, are often co-expressed with fast transmitters in many neuron types ^1-3^. Neuropeptides are packaged into large dense core vesicles (LDCVs), whereas fast transmitters are packaged into synaptic vesicles (SVs) ^3-10^, each capable of different modes of membrane-vesicle fusion ^11^. The release of SVs and LDCVs differ in their response to calcium concentrations; LDCV release typically requires more sustained calcium influx than SV release within a given neuron ^12^. This difference in calcium-dependency results in differing frequency dependencies of SV and LDCV release, with LDCVs typically requiring higher firing frequencies to trigger their release ^6, 13-15^, thus there is preferential SV release at low firing rates, while SV and LDCV co-transmit at higher firing rates. The contents of co-transmitted LDCVs may modulate the function of fast neurotransmitters from a co-expressing neuron, which can shape the gain and temporal resolution of conveyed signals ^16^ and thus afford flexibility and richness to neural output.

However, rapid depletion of SVs during high frequency stimulation has been shown to occur at high release probability synapses, causing transmission of fast neurotransmitters to decrease in fidelity at higher firing frequencies. This would suggest that in specific neural populations, low frequency and high frequency stimulations could result in distinct outputs, predominantly SV and LDCV release respectively, representing a frequency-dependent “switch” in the type of neurotransmitter being released rather than sustained co-release of both at higher frequencies. However, this phenomenon and its behavioral relevance has been difficult to test *in vivo* due to the limited tools available for recording and manipulating SV vs LDCV release.

Previous studies have employed genetic tools to selectively abolish either fast neurotransmitters or neuropeptides from specific neural populations, aiming to discern their respective contributions to behavior ^17-19^. However, the physiological release of SVs and LDCVs in such behavioral contexts has primarily been inferred from measurements of neural activity. Despite these inferences, direct demonstration of SV or LDCV transmission dynamics in behavioral contexts, and their frequency-dependence, has remained elusive due to the lack of tools for monitoring their release *in vivo*.

Here we address this critical gap by directly monitoring and silencing the release of glutamate and neuropeptides from presynaptic terminals in behaving mice. This approach reveals that CGRP^PBel^ neurons switch between the release of SV and LDCVs in response to different frequencies. CGRP^PBel^ neurons respond to appetitive stimuli with low frequency and SV release, whereas aversive stimuli induce high frequency responses accompanied by LDCV release but little SV release. Unexpectedly, we discovered that glutamatergic and peptidergic transmissions from CGRP^PBel^ neurons convey qualitatively distinct signals. Glutamatergic transmission encodes positive valence, while peptidergic transmission encodes negative valence in CGRP^Pbel^ neurons. This discovery reveals a previously unknown capability of CGRP^PBel^ neurons to “switch” between neurotransmitters in a frequency-dependent manner, to bidirectionally regulate positive and negative emotional state.

## Results

### Differential response of CGRP^PBel^ neurons to positive and negative stimulus

Using recently developed genetically encoded presynaptic sensors (Figure 1A) ^20^, we monitored SV and LDCV release from CGRP^PBel^ neuron terminals in the CeAl (the lateral division of the central nucleus of the amygdala). *Calca*^Cre^ mice were unilaterally injected with AAV-DIO-SypSEP (Synaptophysin-Super Ecliptic pHluorin) or AAV-DIO-CybSEP2 (Cytochrome B561-Super Ecliptic pHluorin) in the PBel and had fiberoptic cannulas implanted over the CeAl to detect SV and LDCV release, respectively (Figures 1B,C). A footshock (0.5 mA) preferentially induced increases in CybSEP2 fluorescence over SypSEP fluorescence in the CeAl (Figure 1D). Consistent with what was reported previously during licking of liquid Ensure ^20^, licking of a sucrose solution (10% wt/vol) preferentially induced SypSEP fluorescence rather than CybSEP2 fluorescence (Figure 1E), suggesting that CGRP^PBel^→CeAl terminals preferentially release LDCVs or SVs in response to negative and positive stimuli, respectively, rather than co-transmitting them.

**Figure 1.**
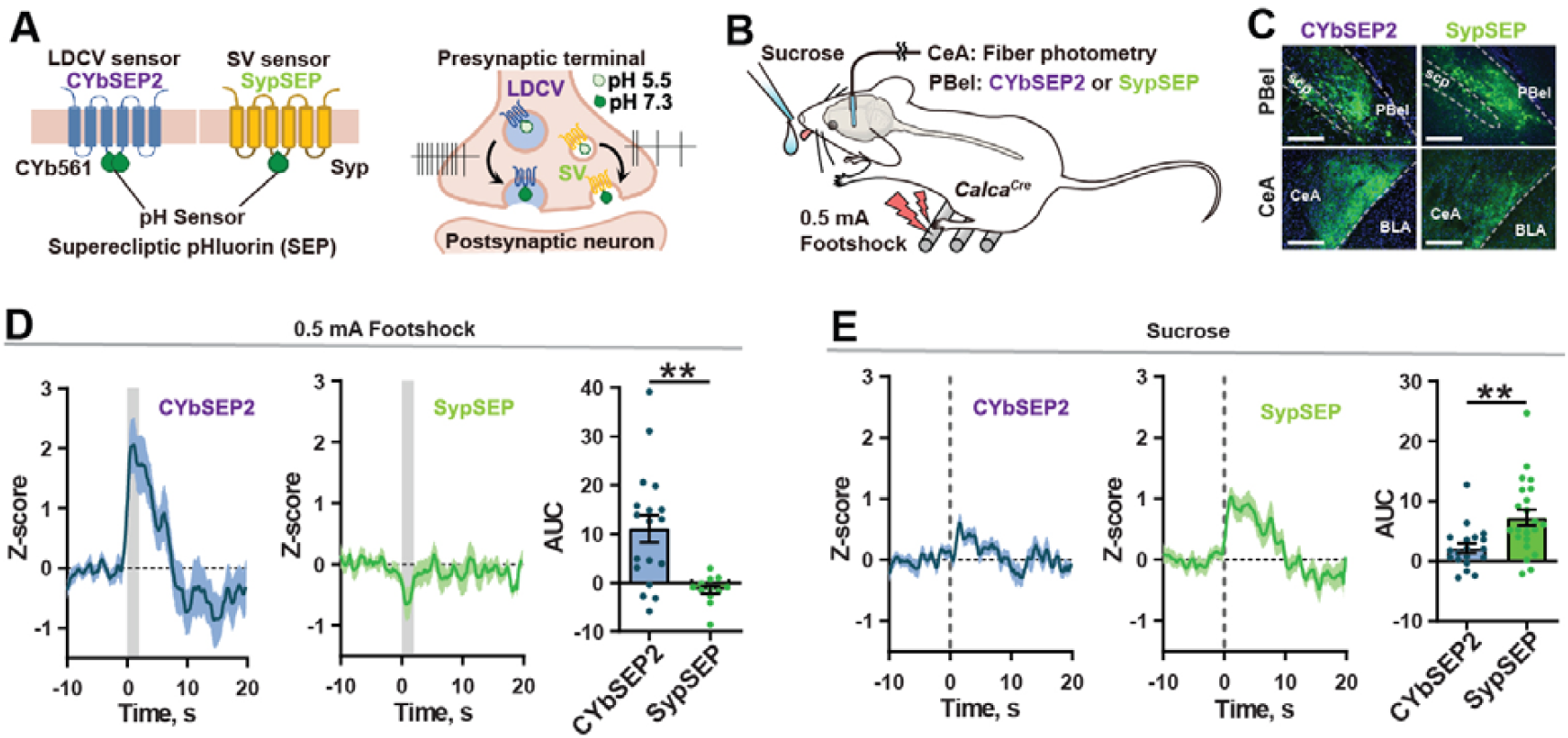
SV and LDCV release from CGRP^PBel^→CeAl terminals in response to positive and negative stimuli. (A) Sensors designed for the detection of LDCV and SV release. (B) LDCV and SV release monitored during presentation of a noxious emotional stimuli. (C) Histological images showing the expression of SV (SypSEP) sensor in the PBel and CeAl. Scale bar, 200μm. (D) Fluorescence signals from CybSEP2 (left) SypSEP (middle) and quantification of AUC data (right) in response to footshock. (E) Fluorescence signals from CybSEP2 (left) SypSEP (middle) and quantification of AUC data (right) in response to sucrose consumption. **P < 0.01. Data are presented as mean ± s.e.m.

To correlate these findings with the activity of CGRP^PBel^ neurons in response to foot shocks and sucrose consumption, we performed fiber photometry recordings. To this end we unilaterally injected *Calca*^Cre^ mice with AAV-DIO-jGCaMP8m and implanted fibreoptic cannulas over the PBel (Figure S1A). Fibre photometry recordings showed that peak calcium responses to foot shock were greater than those seen in response to sucrose consumption (Figures S1B and 1C). Peak amplitudes to each type of stimulus were maintained over repeated stimuli suggesting that CGRP^PBel^ neurons may possibly respond more to valence rather than salience (Figures S1D-G).

Since increased fluorescence in fibre photometry could be interpreted as either with a higher firing frequency or a higher proportion of cells responding, we aimed to distinguish between these possibilities using single-unit recordings. *Calca*^Cre^ mice were unilaterally injected with AAV-DIO-ChR2-eYFP into the PBel and implanted with optetrode drives (Figure 2A). Mice were presented with either a sucrose solution or a tail pinch to evoke positive and negative affect respectively. Among 126 single units recorded in 6 mice, 50 units were further classified as CGRP-positive neurons based on optotagging criteria (Figures S2A-C). Fifty-six percent of the units (28 of 50; 56%) exhibited an increase in firing rate after sucrose consumption (Figures 2B and 2C). All 28 units that responded to sucrose were also activated by tail pinch (Figure 2D). Average changes in firing rate of the “both responsive” units were 8.41 and 37.31 Hz in response to sucrose and tail pinch, respectively (Figure 2E), suggesting that a portion of CGRP^PBel^ neurons can distinguish between positive and negative stimuli via firing frequency. Taken together, these results suggest that positive and negative stimuli lead to preferential release of SVs and LDCVs from CGRP^PBel^ neurons respectively and are associated with differing firing frequencies.

**Figure 2.**
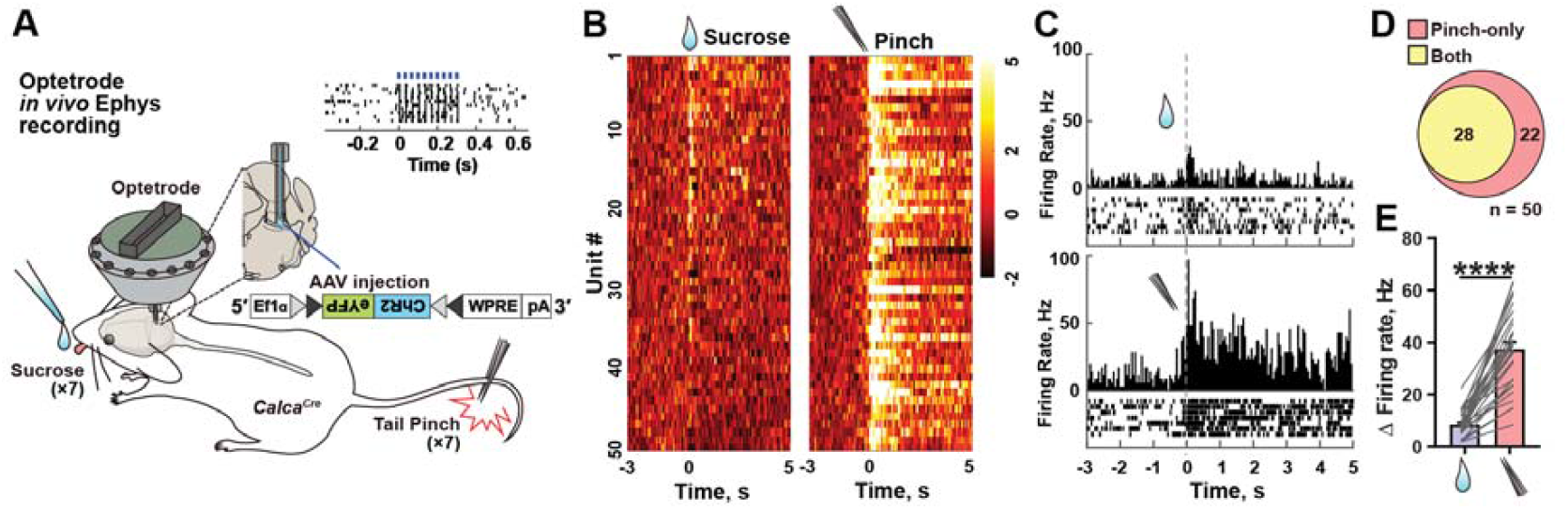
CGRP^PBel^ neurons respond to positive and negative stimuli with differing frequencies. (A) *In vivo* single-unit recording of CGRP^PBel^ neurons using opterode. Right panel (raster plot) shows a light-responsive unit. (B) Z-scored heat map showing normalized firing responses to sucrose and tail pinch. Each row represents the response recorded from the same unit. (C) Firing rate of an example unit responding to both sucrose and tail pinch. (D) Number of optotagged neurons that were activated by pinch-only or both pinch and sucrose. (E) Changes (Δ) in the firing rate induced by sucrose and tail pinch. Baseline firing rate was subtracted from the peri-event firing rate (0-0.5 s). ****P < 0.0001. Data are presented as mean ± s.e.m

### CGRP^PBel^ neurons can bidirectionally induce valence

To validate that firing frequency can mediate the preferential release of SVs or LDCVs, we monitored SV and LDCV release in the CGRP^PBel→CeAl^ terminals while electrically stimulating the PBel. *Calca*^Cre^ mice were implanted with a tungsten electrode and unilaterally injected with AAV-DIO-SypSEP or AAV-DIO-CybSEP2 in the external lateral parabrachial nucleus and had fiberoptic cannulas implanted over the CeAl (Figure 3A). Electrical stimulation of PBel neurons at 5 or 10 Hz evoked little increases in CybSEP2 fluorescence, whereas stimulation at 20 or 50 Hz was able to evoke increases in CybSEP2 fluorescence (Figure 3B). In contrast, SypSEP fluorescence was seen at 5 Hz but began to decrease below baseline levels at 20 Hz stimulation (Figure 3C). To identify the likely fast neurotransmitter being released by CGRP^PBel^ neurons we performed *in situ* hybridization against *Calca* and *Slc17a6* transcripts in the PBel. We found that 95.76 ± 0.65 % of CGRP neurons express Vglut2 (Figures S3A-C). We then recapitulated the results from our SypSEP sensor by unilaterally injecting *Calca*^Cre^ mice with a postsynaptic sensor of glutamate (iGluSnFR) in the CeAl. iGluSnFR fluorescence increased in the CeAl at 2-, 4-, 8-, and 10 Hz electrical stimulation, whereas fluorescence decreased below baseline levels at 20 Hz and 40 Hz stimulation (Figure S4A and 4B). Finally, we performed slice recordings in the CeAl from *Calca*^Cre^ mice injected with AAV-DIO-ChR2-eYFP and AAV-DIO-mCherry in the PBel (Figure 3D). We selectively targeted cells in the CeAl that are directly innervated by the CGRP^PBel^ terminals with perisomatically labelled mCherry signals (Figure 3E). Voltage-clamp recordings showed optically evoked excitatory postsynaptic currents (oEPSCs) at the CeAl, which were abolished in the presence of CNQX, indicating glutamatergic transmission (Figure 3F). oEPSCs displayed less fidelity at 40 Hz compared to 4 Hz, suggesting impaired glutamatergic fidelity at high frequencies (Figures 3G and 3H). Current-clamp recordings of the same CeAl neurons showed slow membrane potential depolarizations during 40 Hz photostimulation of CGRP^PBel^ neuron terminals, whereas this was not seen during 4 Hz photostimulation, suggesting that neuropeptide release from CGRP^PBel→CeAl^ terminals can occur at 40 Hz but not at 4 Hz (Figures 3I and 3J). Optogenetic constructs were validated through slice patch-clamp recordings (Figure S5), where CGRP^PBel^ neurons were able to respond to light pulses reliably at both 4 Hz and 40 Hz. These results suggest that stimulation of the PBel at differential frequencies can lead to release of either SV or LDCVs in the CeAl.

**Figure 3.**
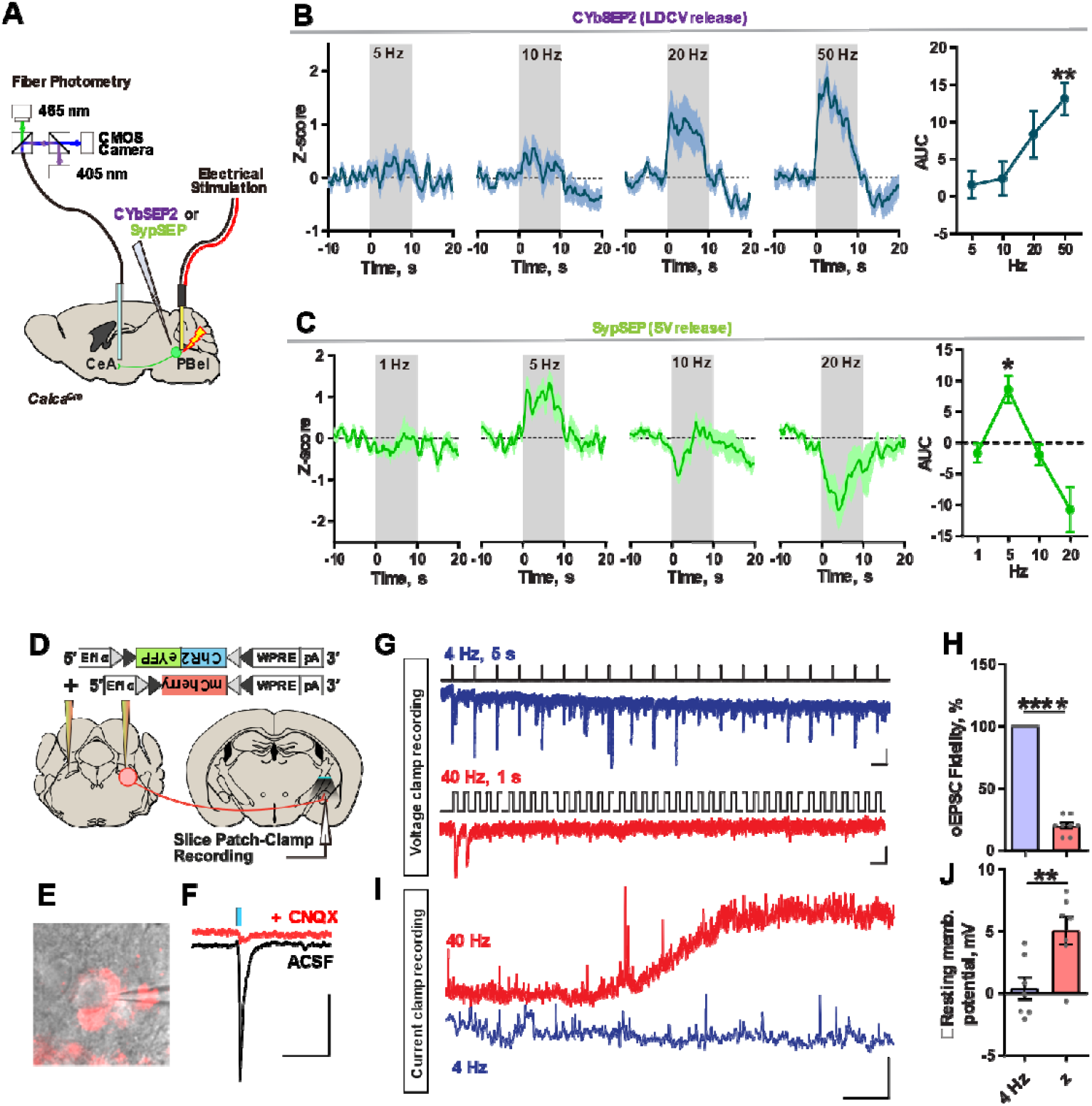
SV and LDCV release from CGRP^PBel^ neurons are driven by differing frequencies. (A) Schematic of monitoring LDCV and SV releases from the CGRP^PBel^→CeAl terminals in vivo. PBel neurons were stimulated using a bipolar tungsten electrode. (B) LDCV release signals during electrical stimulation of the PBel area (left) and quantification of AUC data (right). (N = 4 mice, average of 3-4 repeats). (C) SV release signals during electrical stimulation of the PBel area (left) and quantification of AUC data (right). (N = 4 mice, average of 3 repeats). (D) Schematic of experiment. (E) Image of a patch-clamped CeAl neuron, surrounded by perisomatic mCherry signals. Scale? (F) Optically evoked EPSCs are abolished by CNQX treatment. Scale bar, 50 μs, 50 pA. (G) Post-synaptic current responses evoked by a train of photostimulation at 4 Hz or 40 Hz. Scale bar: 100 ms, 20 pA (top); 20 ms, 20 pA (bottom). (H) Fidelity of oEPSCs in response to 4-Hz and 40-Hz photostimulation. (I) Example trace of a CeAl neurons membrane potential in response to 4-Hz or 40-Hz photostimulation. Scale bar: 30 s, 2 mV. (J) Quantification of membrane potential responses to 4-Hz or 40-Hz photostimulation. ****P < 0.0001, **P < 0.01, *P < 0.05. Data are presented as mean ± s.e.m.

To then establish whether CGRP^PBel^ neurons can differentially drive positive or negative valence according to firing frequency, we performed real-time place preference (RTPP) test. *Calca*^Cre^ mice were bilaterally injected with AAV-DIO-ChR2-eYFP or AAV-DIO-eYFP and bilaterally implanted with fiberoptic cannulas over the PBel (Figure 4A). Mice were placed in a two-chamber box where one side of the chamber was paired with photostimulation (Figure 4B). Pairing one chamber with 40 Hz photostimulation caused ChR2 injected mice to spend less time in that chamber, whereas pairing a chamber with 4 Hz photostimulation caused them to spend more time in that chamber (Figures 4C-E). No differences were seen in eYFP injected mice (Figure S6). These results suggest that optogenetic stimulation of CGRP^PBel^ neurons at differing frequencies can drive opposing valences.

**Figure 4.**
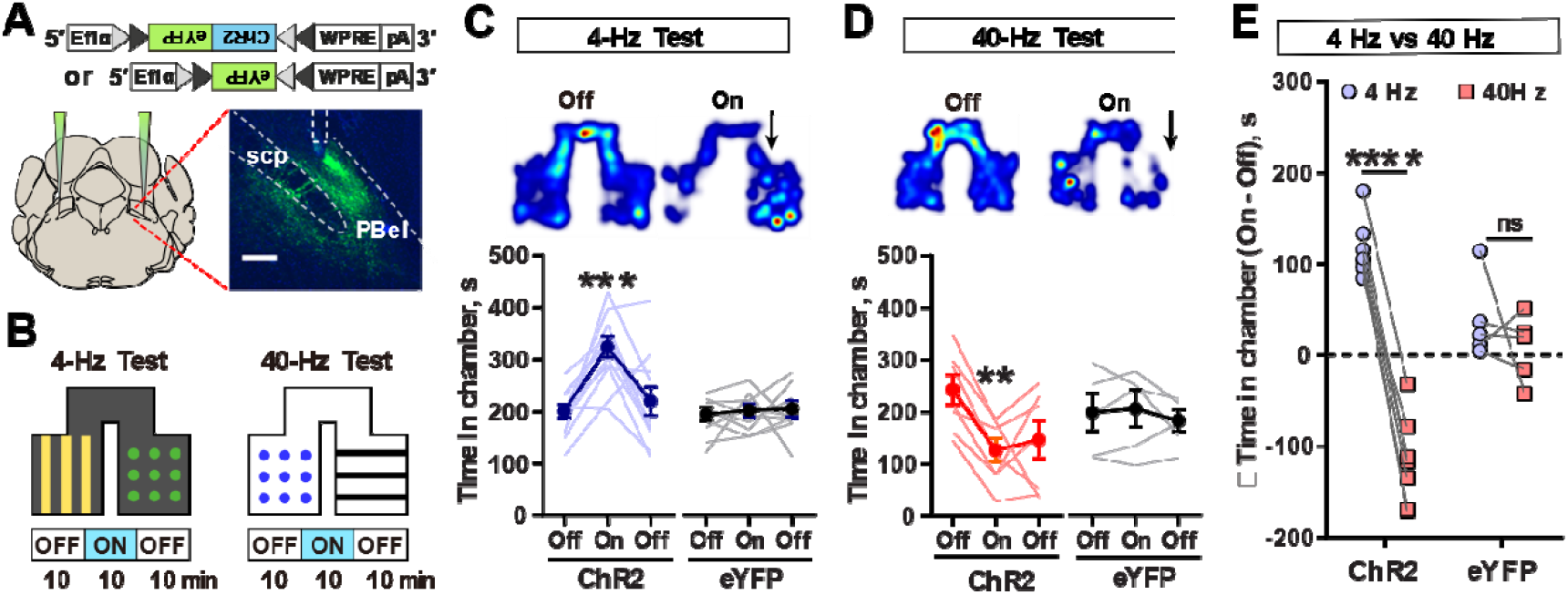
Stimulation of CGRP^PBel^ neurons at differing frequencies induces opposing responses. (A) Schematic of injections. Inset image shows the expression of ChR2-eYFP in CGRP^PBel^ neurons. Scale bar, 200 μm. (B) Schematic of real-time place preference or aversion test (RTPP/RTPA), paired with 4 Hz or 40 Hz stimulation. (C) Heatmap during RTPP using 4 Hz photostimulation (top). Time spent in chamber for ChR2 and eYFP injected animals (bottom). (ChR2 N = 11; eYFP N = 9). (D) Heatmap during RTPA using phasic 40 Hz photostimulation (top). Time spent in chamber for ChR2 and eYFP injected animals (bottom). (ChR2 N = 7; eYFP N = 5). (E) Light-induced changes in time spent in the paired chamber. ****P < 0.0001, ***P < 0.001, **P < 0.01. Data are presented as mean ± s.e.m.

### Glutamate and neuropeptides from CGRP^PBel^ neurons mediate positive and negative valences respectively

To then establish a causal role for neuropeptide and glutamate release in mediating negative and positive valences respectively, we bilaterally injected *Calca*^Cre^ mice in the PBel with AAV-DIO-NEP_LDCV_ (a recently developed LDCV-targeted neuropeptide-specific endopeptidase ^20^), AAV-DIO-sgRNA-*Slc17a6* or AAV-DIO-mCherry, along with AAV-DIO-ChR2-eYFP and then performed RTPP tests. mCherry + ChR2 group spent less time in a chamber paired with 40 Hz photostimulation whereas they spent more time in a chamber paired with 4 Hz photostimulation (Figure 5A). However, NEP_LDCV_ + ChR2 group spent more time in chambers paired with either 4 Hz or 40 Hz photostimulation (Figures 5B and 5C), suggesting that peptidergic transmission is required to mediate negative valence in CGRP^PBel^ neurons.

**Figure 5.**
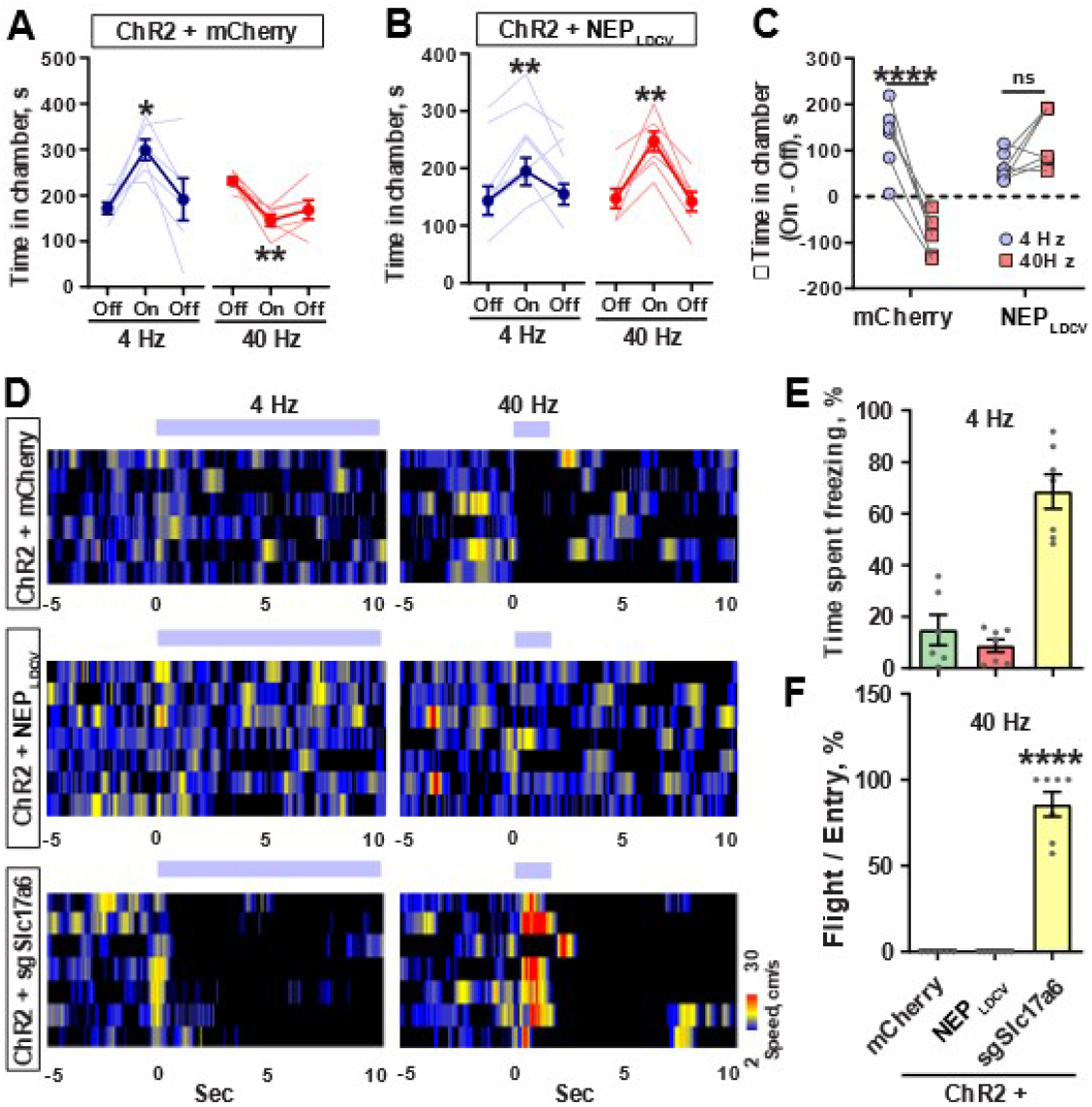
Glutamate and LDCVs from CGRP^PBel^ neurons are required to mediate positive and negative emotional responses, respectively. (A) Time spent in the paired chamber induced by 4 Hz and 40 Hz photostimulation in mCherry injected mice, (N = 6). (B) Time spent in the paired chamber induced by 4 Hz and 40 Hz photostimulation in NEP injected mice, (N = 7). (C) Light-induced change in time spent in the paired chamber induced by 4 Hz or 40 Hz photostimulation. (D) Heatmaps showing the changes in movement speed (cm/s) after the onset of 4 Hz (continuous) and 40 Hz (phasic) photostimulation in mCherry (top), NEP (middle) and sgSlc17A6 (bottom) injected mice. (E) Percentage time spent freezing in chamber paired with 4 Hz photostimulation. (F) Number of flight bouts as a percentage of entries. ****P < 0.0001, **P < 0.01, *P < 0.05. Data are presented as mean ± s.e.m.

Photostimulation of sgRNA-*Slc17a6* + ChR2 group induced behaviours that precluded the use of RTPP tests. Photostimulation at 4 Hz caused profound freezing behaviour, whereas 40 Hz caused immediate flight behaviour in sgRNA-*Slc17a6* + ChR2 group (Figures 5D-F, Figure S7), suggesting that glutamatergic transmission is required to mediate positive valence in CGRP^PBel^ neurons, and activation of CGRP^PBel^ neurons with attenuated glutamatergic transmission exacerbates negative affect. Taken together, these results suggest that neuropeptides release drives negative valence, whereas glutamate release drives positive valence in CGRP^PBel^ neurons.

## Discussion

Here, by direct monitoring and silencing glutamatergic and peptidergic transmissions from the CGRP^PBel^ neurons, we show that glutamate and neuropeptides from CGRP^PBel^ neurons likely encode qualitatively distinct signals. Contrary to our expectations, peptidergic transmission from CGRP^PBel^ neurons did not merely modulate the behavioral output of CGRP^PBel^ neural glutamatergic transmission, but instead induced opposing behavior. These findings are at odds with a previous finding that 5 Hz photostimulation of CGRP^PBel^ neurons can induce conditioned taste aversion ^21^, but we note that this occurred with 2 hours of photostimulation and in dehydrated mice. Given that the lateral parabrachial nucleus is responsive to interoceptive changes, it is possible that the ability of CGRP^PBel^ neurons to encode both negative and positive valence may be influenced by homeostatic deviations and involved in signalling whether incoming stimuli represent deviations away from homeostasis or its restoration.

An alternative hypothesis could be that CGRP^PBel^ neurons release glutamate and neuropeptides in response to stimulus of different intensities. However, we find this explanation to be unlikely given that low frequency and high frequency photostimulation induced preference and aversion respectively. If their release merely reflected stimulus intensity, both would be expected to induce preference or aversion at differing intensities, or be neutral in valence.

Previous studies showed that different transmitters can encode distinct behaviors in sensory neurons in *C. elegans* ^22,23^. However, these studies lack detailed mechanisms explaining how different transmitters can encode distinct information within neurons. Our study addresses this gap by demonstrating the differential release of neurotransmitters in a frequency-dependent manner in neurons, revealing a specific mechanism for how distinct signals are encoded in response to different stimuli. This can be achieved in various ways, such as spatial segregation of the postsynaptic receptors for co-transmitted ligands. This explanation is unlikely to account for our results however, given the presence of CeAl neurons that receive both neuropeptidergic and glutamatergic transmission from CGRP^PBel^ neurons ^*20*^. Instead, our results could be explained due glutamate depletion, leading to peptidergic transmission becoming more dominant. Alternatively, due to the decreasing SypSEP fluorescence seen at higher frequencies, we speculate that another potential mechanism is that SV release is actively inhibited at high frequencies in CGRP^PBel^ neurons. High frequency stimulation has been shown to be able to cause synaptic depression by inhibiting P/Q-type Ca channels through calcium sensor proteins ^24^, impairing further calcium influx and thus impairing further SV release. LDCVs on the other hand are not likely to suffer the same feedback inhibition due to their increased distance from P/Q-type Ca channels, relying primarily more on L-type channels or intracellular calcium stores to trigger release ^25^.

Regardless, the decrease in SypSEP signals suggests that SV release and in turn, SV-based signaling becomes less predominant during high-frequency stimulation. In this model, this functionally allows CGRP^PBel^ neurons to switch from SV to LDCV-based signalling at higher firing frequencies rather than functionally co-signal both continuously and are thus able to convey qualitatively distinct signals by altering from SV to LDCV signaling at higher firing frequencies.

An unexpected pair of results from our experiments include the negative valence seen during 4 Hz photostimulation of CGRP^PBel^ neurons with glutamate abolished and the positive valence seen during 40 Hz photostimulation of CGRP^PBel^ neurons expressing NEP_LDCV_. These results could be interpreted to mean that removal of one neurotransmitter facilitates the release of the other, which we speculate may occur through a competitive mechanism that becomes manifest when the contents of SVs or LDCVs becomes depleted.

Such phenomena can be seen in certain calcium sensing components in vesicular machinery. For example, synaptotagmin-1 is known to be able to mediate spontaneous and evoked synchronous vesicle release, yet its deletion increases spontaneous asynchronous release ^26,27^ which has been interpreted to mean that synchronous and asynchronous modes of vesicle release compete with each other ^27,28^. Thus, it is plausible that a similar competitive mechanism may exist between SV and LDCV release at the level of their calcium-sensing components, but will require future research to test.

Some genetically defined neurons can induce very different types of behaviors depending on context. For example, behaviors associated with mating vs. aggression are encoded by a single population of neurons in the ventromedial hypothalamus that expresses the estrogen receptor type 1 ^29-32^. Early optogenetic studies have also shown that low frequency and high frequency stimulation of defined populations induces distinct, sometimes opposing, behavioral outcomes ^33,34^. Furthermore, past studies have also implicated the lateral parabrachial nucleus in both driving hedonic behaviors ^35,36^, as well as encoding danger signals ^37^, which otherwise seem at odds with each other. Our study provides a potential mechanism for such phenomena, whereby information encoded by glutamate and neuropeptides from such neurons may mediate qualitatively distinct behaviors. The frequency dependency switch between SV and LDCV release, in turn, may give rise to the differing, and sometimes opposing, effects that low frequency vs high frequency stimulation can have. The experimental approach used in this study facilitates the testing of this hypothesis in peptidergic populations outside of CGRP^PBel^ neurons and may reveal differential transmission as a general principle in certain peptidergic circuits of the mammalian brain.

## Acknowledgements

We thank Han lab members for critical discussion of this manuscript. S.H. is supported by 1RF1NS128680 from NINDS, and the Salk Institute Innovation Grant.

## Author Contributions

S.H. conceived of the idea and secured funding. S.H., D-I.K., S.J.K., J.J., Y.S.J., S. P. and M.Y. designed the experiments. D-I.K. performed the transmitter release monitoring experiments and produced AAVs. S.J.K. performed in situ hybridization and calcium imaging experiments. J.J. performed RTPP/RTPA experiments. Y.S.J. and G.H.P. performed and analyzed the optetrode single-unit electrophysiological recording experiment. S.P. and M.Y. performed the slice electrophysiology experiment. S.H., J.J., S.J.K., Y.S.J., S. P. and M.Y. wrote the manuscript.

## Supplementary Materials

Materials and Methods

Figures. S1-S8

Table S1

