## Supplementary information for "Encoding opposing valences through frequency-dependent transmitter switching in single peptidergic neurons"

**The PDF file includes:**

Materials and Methods
Figs. S1 to S8
Table S1

### Materials and Methods

Mice

All protocols for animal experiments were approved by the Institutional Animal Care and Use Committee (IACUC) of the Salk Institute for Biological Studies and Korea University in accordance with the NIH guidelines for animal experimentation. The Calca-Cre transgenic mouse line (expressing Cre-GFP fusion protein in CGRP positive neurons) used in this study was generated by the Richard Palmiter's laboratory (University of Washington Seattle) and backcrossed with C57Bl/6J for > 6 generations. GFP expression from the Cre-GFP fusion protein is undetectable in perfused brain slices, in sharp contrast with the fluorescence from viral expression. Male and female mice were used in all studies. Animals were randomized to experimental groups and no sex differences were noted. Mice were maintained on a normal 12-hour light/dark cycle and provided with food and water ad libitum.

Stereotaxic surgery

Mice (with Calca-Cre transgenic background) were anesthetized with 4% isoflurane and fixed on a stereotaxic surgery frame (David Kopf Instruments). 1.5-2% isoflurane was maintained during the surgery. The cranium was exposed and drilled with a handpiece drill (Foredom) above each injection site. Viral solutions were loaded on a glass pipette filled with mineral oil. Solutions were injected into target brain sites at 1-nL per second. Target coordinates for bilateral PBel were anterior-posterior (AP) -5.1 mm, medial-lateral (ML) ±1.45 mm, dorsal-ventral (DV) -3.7 mm relative to the bregma. Mice were used for behavioral or recording experiments 2-4 weeks after the surgery.

For recording presynaptic release from $CGRP^{PBel}$→CeA by electrical stimulation, AAVDJ-hSyn-DIO-CybSEP2 and AAVDJ-hSYN-DIO-SypSEP were unilaterally injected into the PBel area. A custom-made bipolar electrode was implanted into the PBel. Briefly, electrodes were made of PFA-coated stainless-steel tungsten wires (coated diameter 114.3 µm, A-M systems, Cat#. 791500). Wires were connected to the connector (Dupont connector kit, Plusivo) and then were twisted and cut to 4.5-5 mm in length based on DV value for targeting PBel. Custom-made bipolar electrodes were implanted in the same coordinate for PBel, 15 min after the viral injection. The resistance of generated electrode ranged from 3 to 5 Ω. Custom-made stainless-steel mono fiber-optic cannulas (400 µm, 0.47 NA) were implanted above the CeA (AP: -1.2 mm, ML: -2.85 mm, DV: -4.5 mm). After implantation, superglue was used for fixing the implanted electrodes or optic ferrules followed by dental acrylic to cement onto the skull surface. Behavioral tests were conducted 4 weeks after surgery. For in vivo recording of glutamate release by optogenetics or electrical stimulation, AAV-hSyn-SF-iGluSnFR.A184S (#106174, Addgene) and AAVDJ-DIO-ChrimsonR-mScarlet were unilaterally injected into the CeA and PBel respectively. After viral injection, an optic fiber (200 µm) or bipolar electrode was implanted into the PBel.

For behavioral (RTPP/RTPA) experiments, 300-nL of AAVDJ-EF1a-DIO-hChR2-eYFP-WPRE-pA (1.2E+12 GC/mL) or AAVDJ-hSYN-DIO-eGFP-WPRE-pA (2.3E+12 GC/mL) solution was injected bilaterally into the PBel area. Custom-made ceramic mono fiber-optic cannulas (200 μm, 0.22 NA) were implanted 300 μm above the injection sites and fixed with dental cement. For calcium activity monitoring experiments, 300-nL of AAV1-hSYN-FLEX-jGCaMP8m-WPRE-pA (5E+12 GC/mL) solution was injected unilaterally into the PBel area, followed by the implantation of a custom-made stainless-steel mono fiber-optic cannula (400 μm, 0.37 NA) 200 μm above the injection site. For single-unit recording experiments in vivo, 500 nL of AAV-EF1α-FLEX-ChR2-eYFP-WPRE-pA (7E+12 GC/mL) solution was injected unilaterally into the PBel area followed by the microdrive implantation.

LDCV, SV, and iGluSNFR sensor recording with fiber photometry

CybSEP2, SypSEP, and iGluSNFR sensor responses were recorded using a CMOS sensor-based fiber photometry system (Doric Lenses) operated by a custom-built LabVIEW (National Instrument) software. 465-nm and 405-nm light emission diodes (LEDs) were coupled to dichroic mirrors and then connected to a 20X/0.4 NA objective lens. LED pulses were sinusoidally illuminated at 10 Hz to collect the pH- or glutamate-dependent signals and isosbestic control responses.

For electrical stimulation, mice were attached attached to a fiber-optic patch cord (400 μm, 0.48 NA) which connected to the objective lens of the photometry system, and an electrical patch cord which connects to an electrical stimulator (A-M systems, Model 2100 Isolated Puls Stimulator), and then placed in their home cages for habituation. After habituation for 10 min, 70-100 μA, 1-ms (pulse width) electrical stimulations were given at various frequencies (5 to 40 Hz) for 10 s. Stimulations were repeated for 3-4 times with minimal inter-trial intervals (ITIs) of 10 mins.

For the recording of the response to sucrose and foot shock, mice were placed in an opaque cylinder (11 cm diameter) and habituated for 10 min. After habituation, mice were presented with 10 % of sucrose (10 μL) through a small hole (~1.5 cm diameter) using a micropipette for 3-4 times, and then with an electric foot shock (0.1 or 0.5 mA) for 3-4 repeats. Collected photometry data were analyzed using a custom Python script. Isosbestic control (405 nm; F405) signals were fitted to 465-nm signals (F465) by least mean squares fitting, and the motion corrected fluorescence signals (ΔF/F) were calculated by (F465 – F405fit)/F405fit. ΔF/F was then z-scored relative to the mean and SD of the fluorescence signal and smoothed by 2nd-order Savitzky–Golay filter using Prism 8 (GraphPad software).

Calcium activity monitoring with fiber photometry

Bulk calcium signals from $CGRP^{PBel}$ neurons were monitored using a custom-built fiber photometry system based on the open source pyPhotometry platform (https://pyphotometry.readthedocs.io/en/latest/). A 465 nm LED was used to induce Ca2+-dependent fluorescence signals, and a 405 nm LED was used for Ca2+-independent (isosbestic) fluorescence signals. Analysis was performed in the same way as described in the LDCV and SV sensor recording with fiber photometry section.

Slice electrophysiology recording ex vivo

Mice were anesthetized with isoflurane and transcardially perfused with ice-cold cutting solution (75.0 mM Sucrose, 87.0 mM NaCl, 25.0 mM $NaHCO_3$, 1.25 mM $NaH_2PO_4$, 2.5 mM KCl, 0.5 mM $CaCl_2$, 7.0 mM $MgCl_2$, 25.0 mM glucose, 5.0 mM ascorbic acid and 3.0 mM pyruvic acid, bubbled with 95% $O_2$ and 5% $CO_2$). Mice were decapitated, then brains were quickly removed to ice-cold cutting solution. Coronal slices containing the PBel or CeA (250 µm) were prepared using a VT 1200S Vibratome (Leica) and transferred to a storage chamber containing artificial cerebrospinal fluid (aCSF; 124 mM NaCl, 2.5 mM KCl, 26.2 mM $NaHCO_3$, 1.2 mM $NaH_2PO_4$, 13 mM glucose, 2 mM $MgSO_4$ and 2 mM $CaCl_2$, at 32C, pH 7.4, bubbled with 95% $O_2$ and 5% $CO_2$). After recovery for 30 min at 32°C, slices were transferred to room temperature (22 - 24C) for at least 60 min before use. Slices were transferred into the recording chamber and perfused with aCSF at a flow rate of 2 mL/min. The temperature of the aCSF was controlled at 32°C by a TC-324C Temperature controller (Warner Instruments).

For the single-pulse stimulation recording in CeA slices, for evoked EPSC, a 2-ms blue light pulse was used from a collimated 473 nm light-emitting diode (pE-300, CoolLED) under the control of an Axon Digidata 1440A Data Acquisition System and pClamp 10 software (Molecular Devices). CeA neurons surrounded by the perisomatic mCherry signals (PBel-to-CeA connection) were selected. Each cell was whole-cell clamped with a holding potential at -70 mV with an internal solution (130 mM K-gluconate, 20 mM HEPES, 2 mM NaCl, 4 mM $MgCl_2$, 0.25 mM EGTA, 4 mM Na-ATP, 0.4 mM Na-GTP, pH 7.2). Evoked-EPSCs were blocked by perfusion of 10 mM CNQX (Tocris).

For the EPSC fidelity measurement in CeA slices, 20 light pulses at 4-Hz or 40-Hz frequency were shined onto the cells. For the resting potential measurement, CeA neurons were whole-cell clamped in current-clamp mode and received 4-Hz or 40-Hz light stimulation for 60 s. To confirm ChR2 fidelity, PBel CGRP neurons were optically stimulated at 4-Hz and 40-Hz photostimulation.

Data analysis was performed using the Clampfit module of the pClamp 10 software[JJ1] . Signals were filtered at 10 kHz and digitized at 20 kHz with Digidata 1550B (Molecular Devices). Access resistance was monitored during the recording. If access resistance was larger than 25 MΩ or changed by more than 20%, cell recording was stopped.

Single-unit recording with optotagging in vivo

A microdrive containing one optic fiber (200-µm core diameter, 0.22 NA) and four tetrodes (20 µm-diameter tungsten wire; California Fine Wire) was custom-made. Tetrode tips were cut to be about 400-500 µm longer than the fiber tip and were gold-plated to reach impedances of 200-500 kΩ tested at 1 kHz. After 2 weeks of recovery from the surgery, mice were placed in a holding cage and single-unit activity was monitored using a Cheetah data acquisition system (Digital Lynx SX, Neuralynx). Neuronal signals were filtered between 0.6 and 6 kHz, digitized at 32 kHz, and amplified 1000-8000 times. When unit spikes exceeded a predetermined threshold, the waveforms were recorded for 1 ms. To screen the presence of ChR2-expressing units, 10 blue light pulses (473 nm; 5-ms width; 4-15 mW/mm2 intensity; Laserglow technologies) were provided at 30 Hz via the optic fiber. If no light-responsive units were found, all tetrodes were moved down by 80 μm increments. Once light-responsive units were first found, the animal was habituated to the behavioral recording procedures for 3 consecutive days in which several 10% sucrose (10 μl) and tail pinches (0.5 s) were given in different contexts (sucrose in a white Plexiglass cylinder with a hole for access to sucrose; tail pinches in a translucent rectangle Plexiglass cage). The order of two stimuli was randomized on each day. Then the actual behavioral recording sessions started. On the testing day, spontaneous spikes from PBN neurons were recorded for 10 min in the animal's home cage. The neuronal responses to the sucrose and foot shock (7 trials per stimulus, with inter-trial intervals of 60-90 s) were further recorded. At the end of the session, the neuronal discharge patterns in response to 10 trains of 10 light pulses (total 100 presentations; 30-s inter-train interval) were recorded for identifying ChR2-expressing neurons (opototagging). After the daily recording session, all tetrodes were lowered in 40-80 μm to find other light-responsive neurons and the mouse was returned to its home cage.

To analyze the unit data, unit spikes were isolated based on the various waveform features using Offline Sorter (Plexon). Only units with stable firing activities throughout the behavioral recording sessions were further analyzed using Matlab software (MathWorks). To categorize *Calca* (CGRP+) neurons, a cluster analysis was performed based upon spike probability and latency in response to 100 light pulses. The cluster that had a spike probability of > 0.83 and latency of < 6 ms was classified as *Calca* neurons. These neurons also showed the highest correlation between spontaneous and light-evoked waveforms. To further examine the *Calca* neuronal responses to given stimuli, peri-event time histograms (PETHs; 50-ms bins) were constructed around the stimulus onsets. A *Calca* neuron was considered as responsive to each stimulus if its firing rate during a 0.5-s epoch from the stimulus onset (-0.05 to 0.45 s) was significantly different from the baseline firing rate before the stimulus (-3 to -0.2 s; Wilcoxon signed-rank test).

Real-time place preference/aversion (RTPP/RTPA) test

Mice were connected to an optical patch cord, and placed in a two-chamber testing arena (left/right chamber dimensions with 20 × 25 × 40 cm, W × D × H; two chambers were connected by a neutral bridge area 5 × 20 × 40 cm), and allowed to freely explore the environment for 30 min. Behavior was video-recorded from the top, and the location (body center) of mice was monitored using EthoVision XT (Noldus) software in real-time. One of the two chambers (counterbalanced, for each mouse) was paired with the 4-Hz or 40-Hz light stimulation during the second epoch (10-20 min) of the test. Time spent in the stimulated chamber, bridge area, and opposite chamber was automatically calculated by the EthoVision XT software.

A 470-nm collimated diode laser was used for the photoactivation experiments. Square-pulse TTL signals for controlling the laser device was generated by Pulse Pal device. During behavioral tests, bilateral fiber-optic cannulas were connected to the laser device through an optical patch cord. 4-Hz (5-ms pulse width, ~5 mW power) or 40-Hz phasic (5-ms pulse width, ~5 mW power; 2-s ON and 10-s OFF cycles were repeated) stimulation was delivered during the second epoch.

Histology

Mice were sacrificed by transcardial perfusion with phosphate-buffered saline (PBS) then with 4% paraformaldehyde (PFA; dissolved in phosphate buffer, PB). Dissected brains were incubated in 4% PFA/PB for 16 hours and dehydrated in 30% sucrose for 48 hours. Brains were sectioned into 50-μm coronal slices using a cryotome (-20°C, Leica Biosystems). Brain sections were mounted on microscopic slides and cover-slipped with DAPI Fluoromount-G mounting media (Vectashield Hard Set mounting medium, Vector Laboratories). Images were taken with a BZ-X710 fluorescence microscope (Keyence) for histological observation.

In situ hybridization

Following rapid decapitation of WT mice, brains were frozen in liquid nitrogen with OCT compound and stored at -80° C. Coronal sections containing PBel regions were cut in 10 μm-thick slices at -20°C and thaw-mounted onto slides. in situ hybridization was performed according to the RNAScope 2.0 Fluorescent Multiple Kit User Manual for Fresh Frozen Tissue (Advanced Cell Diagnostics, Inc., USA). Slides containing the specified coronal brain slices were fixed in 4% paraformaldehyde, dehydrated, and pretreated with protease IV solution for 30 min. Slices were then incubated with either target probes for Mm-Calca (RNAScope Probe #578771, Advanced Cell Diagnostics), Mm-Slc17a6 (RNAScope probe #319171-C3, Advanced Cell Diagnostics) for 2 hr. Following probe hybridization, slices underwent a series of probe signal amplification steps (AMP1–4). Slides were cover-slipped with DAPI Fluoromount-G mounting media (Vectashield Hard Set mounting medium, Vector Laboratories). Images were taken with a BZ-X710 fluorescence microscope (Keyence) for histological observation.

Statistics

Statistical analyses were performed using GraphPad Prism 6 software. Parametric tests were used if the normality and equivariance assumptions were satisfied, otherwise non-parametric tests were used for the analysis. Normal distribution of datasets was tested by Shapiro-Wilk normality test ($n \geq 7$) or Kolmogorov-Smirnov test ($n < 7$). Details of statistical tests and results are described in the figure legend and supplementary statistics table.

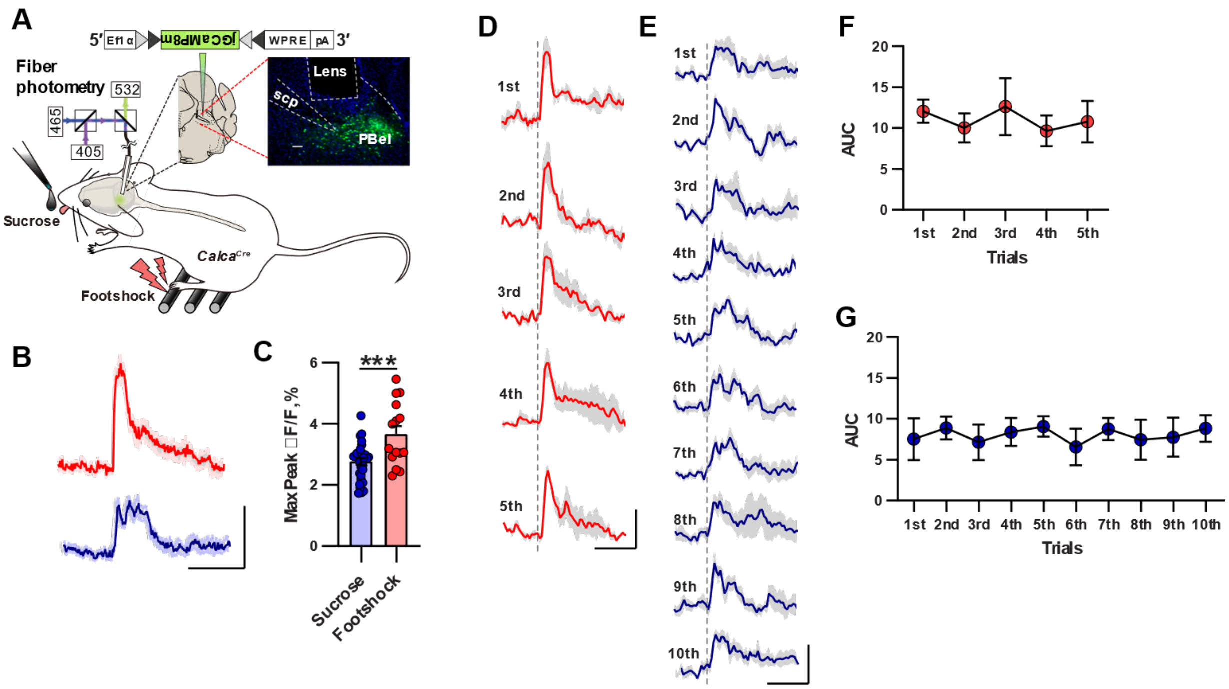


**Fig. S1. CGRP$^{PBel}$ activity during footshocks and sucrose consumption.**

(A) Schematic of fiber photometry and histological image. Scale bar, 100 µm.

(B) Averaged calcium responses to foot shock (red) and sucrose consumption (blue). Scale bar: 2 s and 2 % dF/F. (n = 3 mice).

(C) Peak amplitude of calcium responses for each episode of stimulus presentation (Sucrose, n = 30 episodes; Foot shock n = 15 episodes).

(D Averaged calcium responses to consecutive foot shocks. Scale bar: 2 s and 2 % dF/F.

(E) Averaged calcium responses to consecutive bouts of sucrose consumption. Scale bar: 2 s and2 % dF/F.

(H) Quantification of AUC for data in (E)

(F) Quantification of AUC for data in (D)

***P < 0.001. Data are presented as mean ± s.e.m

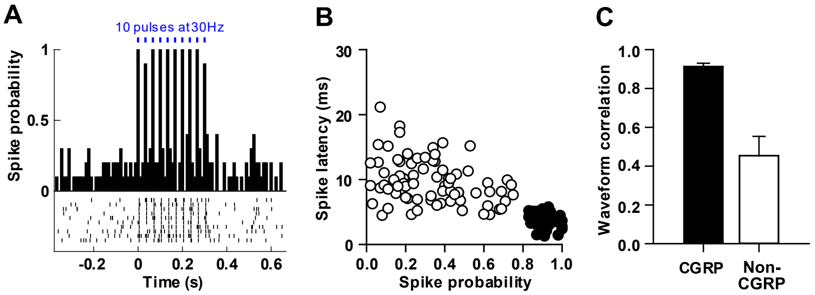


**Fig. S2. Spiking properties of optotagged CGRP units.**

(A) Example of an optotagged CGRP unit. Averaged spike probability (Top). Raster plot of firing responses (bottom).

(B) Distinct properties (spike latency and probability) of neurons classified as CGRP-positive (black) and non-CGRP (white).

(C) Waveform correlation of CGRP-positive and non-CGRP units.

Data are presented as mean ± s.e.m.

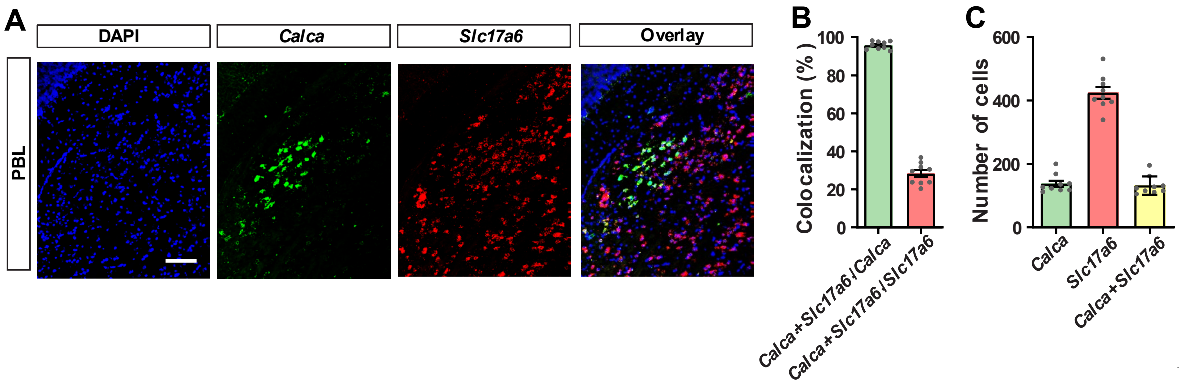


Fig.

**S3. Colocalization of *Calca* and *Slc17a6* mRNAs in the PBel**

(A) Images showing colocalization between *Calca* and *Slc17a6* mRNA signals in a coronal section of the PBel (AP -5.1 mm from the bregma). Scale bar, 100 µm.

(B) quantification of colocalization between *Calca* and *Slc17a6* mRNA.

(C) Total number of *Calca*—positive and *Slc17a6*-positive neurons in the PBel.

Data are presented as mean ± s.e.m.

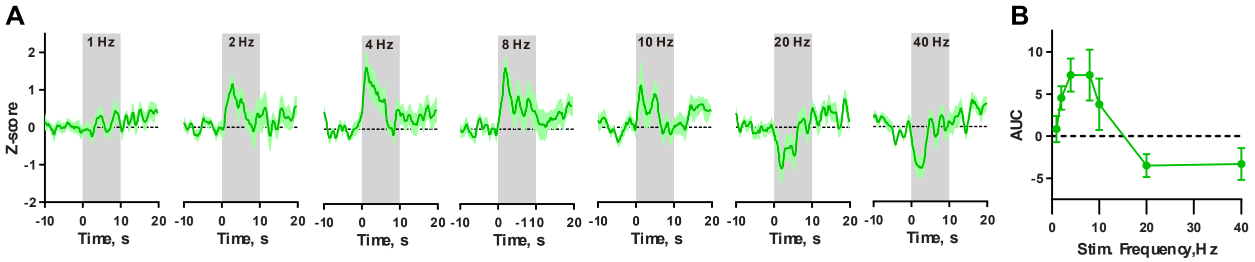


**Fig. S4. iGluSnFR recordings from the CeA at different stimulation frequencies.**

(A) Postsynaptic iGluSnFR signals in the CeA in response to electrical stimulation of the PBel. (n = 3 mice).

(B) Quantification of AUC data in (A).

Data are presented as mean ± s.e.m.

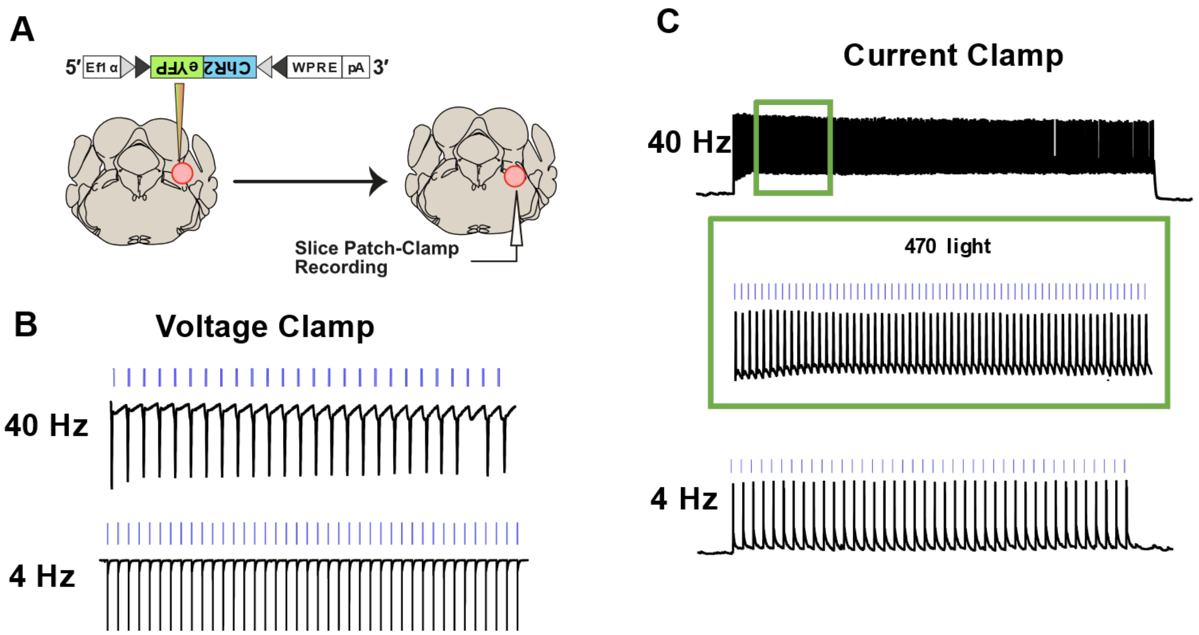


**Fig. S5. CGRP$^{PBel}$ neurons reliably respond to soma photostimulation**

(A) Schematic of experiment CGRP$^{PBel}$ neurons were whole-cell patch clamped and recorded during photostimulation.

(B) Representative traces of a cell showing a current response to 4-Hz and 40-Hz photostimulation.

(C) Representative traces of a cell showing membrane potential responses to 4-Hz and 40-Hz photostimulation.

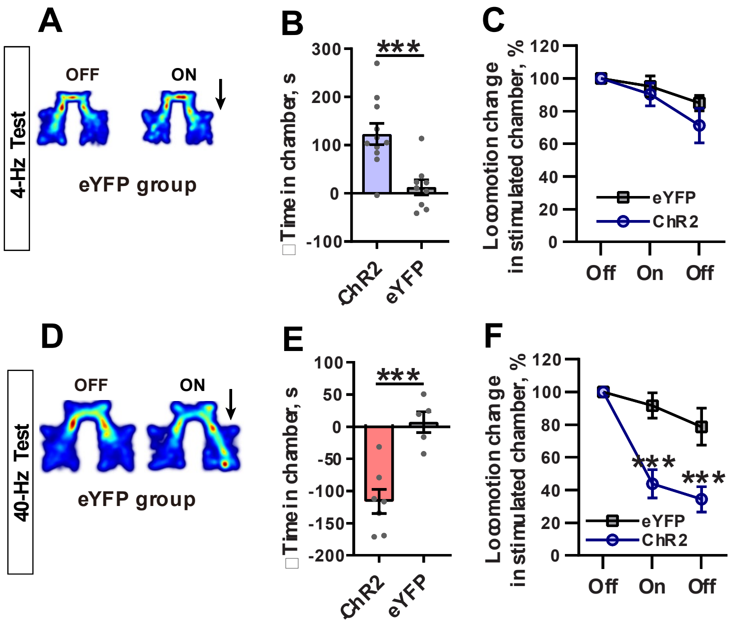


**Fig. S6. Photoactivation of CGRP$^{PBel}$ neurons in eYFP-injected mice does not induce emotional responses**

(A) Representative heatmap of EYFP injected mouse.

(B) Light-induced change in time spent in the paired chamber, induced by 4-Hz photostimulation.

(C) Change in locomotor activity of mice in 4-Hz photostimulation paired chamber.

(D) Representative heatmap of EYFP injected mouse.

(E) Light-induced change in time spent in the paired chamber, induced by 40-Hz photostimulation.

(F) Change in locomotor activity of mice in 40-Hz photostimulation paired chamber

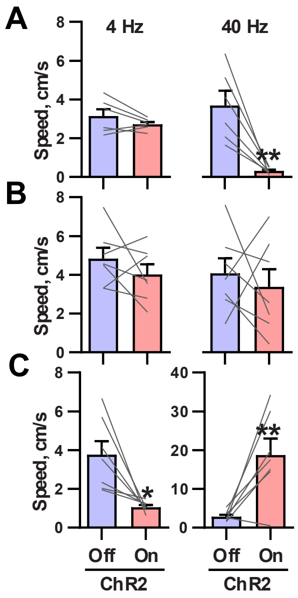


**Fig. S7. Locomotor speed of mice during RTPP/RTPA**

(A) Locomotor speed of mCherry injected mice under 4-Hz (left) and 40-Hz (right) photostimulation.

(B), Locomotor speed of NEPLDCV injected mice under 4-Hz (left) and 40-Hz (right) photostimulation.

(C) Locomotor speed of sg*Slc17a6* injected mice under 4-Hz (left) and 40-Hz (right) photostimulation.

*P < 0.05, **P < 0.01. Data are presented as mean ± s.e.m.

### Table S1. Supplementary statistics table

Supplementary statistical information for each figure

| Figure | Experiment | Cohort | Compared effects | N | P | test |
|---|---|---|---|---|---|---|
| Fig 1D | PBel-CeA CYbSEP2/ SypSEP activity by footshock | PBel Calca CYbSEP2/ SypSEP | AUC by footshock | 18/ 5 and 12 trials/ 4 mice | t = 3.547, df = 28<br>P = 0.0014 | Unpaired t test, Two tailed |
| Fig 1E | PBel-CeA CYbSEP2/ SypSEP activity by sucrose | PBel Calca CYbSEP2/ SypSEP | AUC by sucrose | 19/ 5 and 23 trials/ 5 mice | t = 3.178, df = 40<br>P = 0.0029 | Unpaired t test, Two tailed |
| Fig 2E | PBel *in-vivo* electrophysiology during sucrose vs pinch | PBL Calca ChR2 | Firing rate change by sucrose/ pinch | 28 cells/ 6 mice | t = 10.79, d f= 27<br>P < 0.0001 | Paired t test, Two tailed |
| Fig 3B | PBel-CeA CYbSEP2 activity by electrical stimulation | PBel Calca CYbSEP2 | AUC by different frequencies | 14 trials/ 4 mice | F (2.408, 31.30) = 6.410, P = 0.0030<br>5-Hz vs 50-Hz, P = 0.0012 | RM 1-way ANOVA Sidak's |
| Fig 3C | PBel-CeA SypSEP activity by electrical stimulation | PBel Calca SypSEP | AUC by different frequencies | 12 trials/ 4 mice | F (2.142, 23.56) = 10.30, P = 0.0005<br>1-Hz vs 5-Hz, P = 0.0108<br>1-Hz vs 20-Hz, P = 0.0483 | RM 1-way ANOVA Sidak's |
| Fig 3H | 4-Hz vs 40-Hz light evoked PBel-CeA EPSC | PBel Calca ChR2 | oEPSC fidelity | 8 cells/ 5 mice | t = 29.93, df = 7<br>P < 0.0001 | Paired t test, Two tailed |
| Fig 3J | 4-Hz vs 40-Hz light induced CeA neuron resting membrane potential | PBel Calca ChR2 | Difference of resting membrane potential | 7 cells/ 5 mice | t = 5.676, df = 6<br>P = 0.0013 | Paired t test, Two tailed |
| Fig 4C | PBel 4-Hz activation in RTPP | PBel Calca ChR2/ eYFP | Time in chamber by laser | 11 and 9 mice | ChR2:<br>F (1.566, 15.66) = 10.15, P = 0.0025<br>Off1 vs On: P = 0.0004<br><br>EYFP:<br>F (1.432, 11.46) = 0.1711, P = 0.7734 | RM 1-way ANOVA Sidak's |
| Fig 4D | PBel 40-Hz activation in RTPA | PBel Calca ChR2/ eYFP | Time in chamber by laser | 7 and 5 mice | ChR2:<br>F (1.138, 6.829) = 5.584, P = 0.0483<br>Off1 vs On: P = 0.0016<br><br>EYFP:<br>F (1.239, 4.955) = 0.4153, P = 0.5904 | RM 1-way ANOVA Sidak's |
| Fig 4E | PBel 4-Hz vs 40-Hz activation in RTPP/RTPA | PBel Calca ChR2/ eYFP | Difference in time in chamber by laser | 7 and 5 mice | F (1,10) = 39.36, P < 0.0001<br>ChR2: 4-Hz vs 40-Hz, P < 0.0001 | RM 2-way ANOVA Sidak's |
| Fig 5A | PBel 4-Hz vs 40-Hz activation in RTPP/RTPA | PBel Calca ChR2 + mCherry | Time in chamber by laser | 6 mice | 4-Hz:<br>F (1.659, 8.297) = 6.743, P = 0.0212<br>Off1 vs On: P = 0.0163<br><br>40-Hz:<br>F (1.400, 7.002) = 9.939, P = 0.0122<br>Off1 vs On: P = 0.0083 | RM 1-way ANOVA Sidak's |
| Fig 5B | PBel 4-Hz vs 40-Hz activation in RTPP/RTPA | PBel Calca ChR2 + NEP | Time in chamber by laser | 7 mice | 4-Hz:<br>F (1.273, 7.640) = 4.441, P = 0.0640<br>Off1 vs On: P = 0.0026<br><br>40-Hz:<br>F (1.446, 8.676) = 30.88, P = 0.0002<br>Off1 vs On: P = 0.0040 | RM 1-way ANOVA Sidak's |
| Fig 5C | PBel 4-Hz vs 40-Hz activation in RTPP/RTPA | PBel Calca ChR2 + mCherry/ NEP | Difference of time in chamber | 6 and 7 mice | F (1,11) = 55.98, P < 0.0001<br>ChR2+mcherry: 4-Hz vs 40-Hz, P < 0.0001<br>4Hz: ChR2 + mCherry vs ChR2 + NEP, P < 0.0001 | RM 2-way ANOVA Sidak's |
| Fig 5E | Freezing behavior by PBel 4-Hz activation | PBel Calca ChR2 + mCherry/ NEP/ Vglut2 KO | Freezing time by 4-Hz laser | 6, 7 and 7 mice | F (2, 17) = 39.69, P < 0.0001<br>ChR2 + mCherry vs ChR2 + Vglut2 KO, P < 0.0001<br>ChR2 + NEP vs ChR2 + Vglut2 KO, P < 0.0001 | RM 1-way ANOVA Sidak's |
| Fig 5F | Flight/entry behavior by PBel 40-Hz activation | PBel Calca ChR2 + mCherry/ NEP/ Vglut2 KO | Flight/Entry by 40-Hz laser | 6, 7 and 7 mice | F (2, 17) = 130.8, P < 0.0001<br>ChR2 + mCherry vs ChR2 + Vglut2 KO, P < 0.0001<br>ChR2 + NEP vs ChR2 + Vglut2 KO, P < 0.0001 | RM 1-way ANOVA Sidak's |
| fig. S1C | PBel *in-vivo* calcium activity during sucrose vs footshock | PBel Calca jGCaMP8m | Max peak by sucrose/ footshock | 30/3 and 15 trials/ 3 mice | t = 29.93, df = 7<br>P < 0.0001 | Unpaired t test, Two tailed |
| fig. S2C | PBel *in-vivo* electrophysiology during sucrose vs pinch | PBel CGRP/ Non-CGRP | Waveform correlation | 50 and 76 cells/ 6 mice | t = 3.724, df = 124<br>P = 0.0003 | Unpaired t test, Two tailed |
| fig. S3B | RNAscope with *Calca* and *Slc17a6* in WT PBel | PBel *Calca*/ *Slc17a6* | Colocalization | 9 slices/ 3 mice | t = 34.66, df = 8<br>P < 0.0001 | Paired t test, Two tailed |

| | | | | | | |
|---|---|---|---|---|---|---|
| fig. S3C | RNAscope with *Calca* and *Slc17a6* in WT PBel | PBel *Calca*/ *Slc17a6*/ *Calca+Slc17a6* | Number of cells | 9 slices/ 3 mice | F (1.007, 8.055) = 568.2, P < 0.0001<br>*Calca* vs *Slc17a6*, P < 0.0001<br>*Slc17a6* vs *Calca* + *Slc17a6*, P < 0.0001 | RM 1-way ANOVA Sidak's |
| fig. S4B | PBel-CeA igluSnRF activity by electrical stimulation | PBel electrode + CeA igluSnRF | AUC by different frequency | 14 trials/ 3 mice | F (3.559, 46.26) = 4.554, P = 0.0047 | RM 1-way ANOVA Sidak's |
| fig. S6B | PBel 4-Hz activation in RTPP | PBel Calca ChR2 / eYFP | Difference of time in chamber | 11 and 9 mice | t = 3.939, df = 18<br>P = 0.001 | Unpaired t test, Two tailed |
| fig. S6C | PBel 4-Hz activation in RTPP | PBel Calca ChR2 / eYFP | Locomotion changes in stimulated chamber | 11 and 9 mice | F (2, 36) = 0.7274, P = 0.4901 | RM 2-way ANOVA Sidak's |
| fig. S6E | PBel 40-Hz activation in RTPA | PBel Calca ChR2 / eYFP | Difference of time in chamber | 7 and 5 mice | t = 4.711, df = 10<br>P = 0.0008 | Unpaired t test, Two tailed |
| fig. S6F | PBel 40-Hz activation in RTPA | PBel Calca ChR2 / eYFP | Locomotion changes in stimulated chamber | 7 and 5 mice | F (2, 20) = 8.536, P = 0.0021<br>On: ChR2 vs EYFP, P = 0.0002<br>Off2: ChR2 vs EYFP, P = 0.0005 | RM 2-way ANOVA Sidak's |
| fig. S7A | PBel 4-Hz vs 40-Hz activation in RTPP/RTPA | PBel Calca ChR2 + mCherry | Average speed by laser | 6 mice | 4-Hz: t = 1.521, df = 5, P = 0.1888<br>40-Hz: t = 4.514, df = 5, P = 0.0063 | Paired t test, Two tailed |
| fig. S7B | PBel 4-Hz vs 40-Hz activation in RTPP/RTPA | PBel Calca ChR2 + NEP | Average speed by laser | 7 mice | 4-Hz: t = 1.137, df = 6, P = 0.2988<br>40-Hz: t = 6.026, df = 6, P = 0.6026 | Paired t test, Two tailed |
| fig. S7C | PBel 4-Hz vs 40-Hz activation in RTPP/RTPA | PBel Calca ChR2 + Vglut2 KO | Average speed by laser | 7 mice | 4-Hz: t = 3.391, df = 6, P = 0.0147<br>40-Hz: t = 3.729, df = 6, P = 0.0097 | Paired t test, Two tailed |
